# An enzyme-level benchmark based on environmental bacterial laccases for predicting contaminant fate in water

**DOI:** 10.64898/2026.01.27.701970

**Authors:** Yaochun Yu, Kunyang Zhang, Vincenz-Maria Steiner, Victoria Poltorak, Silke I. Probst, Serina L. Robinson, Jürg Hutter, Hiroko Satoh, Kathrin Fenner

## Abstract

Bacterial laccases are widespread multicopper oxidases whose roles in the fate of anthropogenic chemicals in aquatic environments remain poorly understood. Here, we integrate metagenomic analysis, a miniaturized high-throughput assay and machine learning to establish an enzyme-level benchmark for predicting biotransformation of wastewater-relevant trace organic contaminants by laccase-mediator systems. Using a laccase from an ammonia-oxidizing bacterium as a model enzyme, we screened 183 compounds and identified 38 that underwent significant removal. Following phylogenetic analysis of environmental homologs, we expressed and purified two additional laccases from the bacterial methanotrophic phylum *Methylmirabilota* and an archaeal phylum *Thermoproteota*, demonstrating the activity of this enzyme family across domains. Graph convolutional network models trained on the dataset achieved up to 78% accuracy in classifying degradable versus persistent chemicals, while quantum-chemical descriptors highlighted key electronic properties governing oxidation. This bottom-up approach to enzyme-chemical interactions establishes a trajectory towards predicting contaminant persistence in engineered and natural waters.

## Introduction

The European Commission’s Safe and Sustainable by Design (SSbD) Framework aims to integrate safety and sustainability considerations at the earliest stages of chemical innovation, rather than addressing them afterwards^1^. A central component of this effort is the ability to anticipate how new chemicals will behave once they enter the environment, for example, estimating their environmental half-lives^2^, identifying potential biotransformation pathways^3^, and assessing whether their transformation products pose additional risks^4, 5^. Yet, despite substantial research on contaminant persistence, the mechanistic understanding of many biotransformation reactions remains limited^6^.

The knowledge gap persists for several reasons. Many studies still emphasize system-level parent-compound removal rather than the underlying biochemical mechanisms, leaving the enzymes responsible for individual reactions largely unresolved. Moreover, because many environmentally relevant microorganisms remain uncultivated, direct validation of their metabolic capabilities is limited^7, 8^. Although meta-omics sequencing and high-resolution mass spectrometry can track community shifts and detect transformation products, they rarely provide direct evidence needed to attribute specific biotransformation steps to particular microbial taxa or enzymes^9^. Biochemical approaches, including metabolomics-guided bioinformatics and *in vitro* functional assays, can validate the enzyme activities^6^, but available data remain insufficient to elucidate reactivity across diverse contaminant classes. Enzymes can be highly promiscuous^6, 10^, and without understanding this promiscuity across environmental homologs, linking chemical structure to community-encoded function, and ultimately predicting chemical fate from genomic information, remains difficult.

Nowadays, artificial intelligence (AI) is beginning to show promise for predicting chemical properties and ecological impacts^11–13^. For example, recent work has used wastewater treatment data to predict contaminant removal efficiency^14–18^, and physicochemical parameters such as lipophilicity data to predict toxicity toward nitrifying microorganisms^19–22^. However, the use of AI in this context is still strongly limited by the type and quality of data that are available^23, 24^. From a chemical perspective, predictive models require reactivity data generated under well-defined and comparable conditions, whereas existing datasets are often sparse, inconsistent or difficult to compare across studies. From a microbial perspective, metagenomic and metatranscriptomic datasets contain rich community-level genetic information, yet the specific functional determinants of individual biotransformation steps are often difficult to extract for predictive modeling. Enzyme activity provides a complementary layer that helps link community-scale sequence information to chemical biotransformation outcomes. On this basis, we hypothesize that a bottom-up strategy focused on enzyme–chemical reactivity provides a practical starting point for reducing system complexity while retaining the resolution required for chemical fate prediction.

In this study, we focused on environmental bacterial laccases, a widespread family of multicopper oxidases, as a model system to test this hypothesis. Using a previously identified bacterial laccase from activated sludge as a reference, we performed metagenomic analyses to map its global distribution, identify more than one thousand environmental homologs, and confirm oxidation activity in two additional phylogenetically distinct representatives. We then applied a laccase–mediator system to create a chemically well-defined oxidative environment and used a miniaturized high-throughput assay to measure the reactivity of 183 wastewater-relevant trace organic contaminants under standardized conditions^25, 26^. To support downstream prediction, we developed a data-cleaning strategy that produces a consistent and machine-learning-ready reactivity dataset. We further trained graph convolutional network models to classify reactive versus persistent chemicals and used quantum chemical descriptors to interpret the key electronic features governing oxidation. Collectively, this study establishes an enzyme-level reactivity benchmark that links biochemical data with predictive modeling, offering a foundational step toward SSbD-aligned tools for assessing the environmental fate of emerging contaminants.

## Main text

### Prevalence of microorganisms encoding homologs of a novel bacterial multicopper oxidase (MCO1) in global environments

In our previous study, we characterized a novel bacterial multicopper oxidase (MCO1) from *Nitrosomonas* sp., identified through metatranscriptomic sequencing, and demonstrated its capability to degrade eight trace organic contaminants (TrOCs) commonly found in wastewater^26^. However, the environmental prevalence and phylogenetic breadth of this enzyme family remained unclear. Here, we assessed the global distribution and taxonomic diversity of MCO1 homologs using four databases (the National Center for Biotechnology Information (NCBI) non-redundant protein sequences database, the Global Microbial Gene Catalog^27^, the BV-BRC database^28^, and the Joint Genome Institute Integrated Microbial Genomes (JGI-IMG) database^29^. Our biogeographic analysis revealed that MCO1 homologs were detected in environmental samples spanning at least five continents (Figure 1, Table S1). These homologs were found predominantly in the genomes of bacteria from soil, marine and freshwater systems, lakes, sediments, and the hypersaline Ekho Lake in Antarctica^30^. Genomes of bacterial strains encoding MCO1 homologs have also been isolated from contaminated environments including a crude-oil contaminated seashore^31^ and soil polluted with chromate^32^ (Figure 1), consistent with previous reports of multicopper oxidases involved in hydrocarbon oxidation^33^ and chromate detoxification^34^. While the presence of MCO1 homologs does not by itself establish a direct role in contaminant biotransformation, it is consistent with the broader observation that multicopper oxidases are detected across diverse environments and can participate in redox processes relevant to contaminant transformation^35–37^.

**Figure 1.**
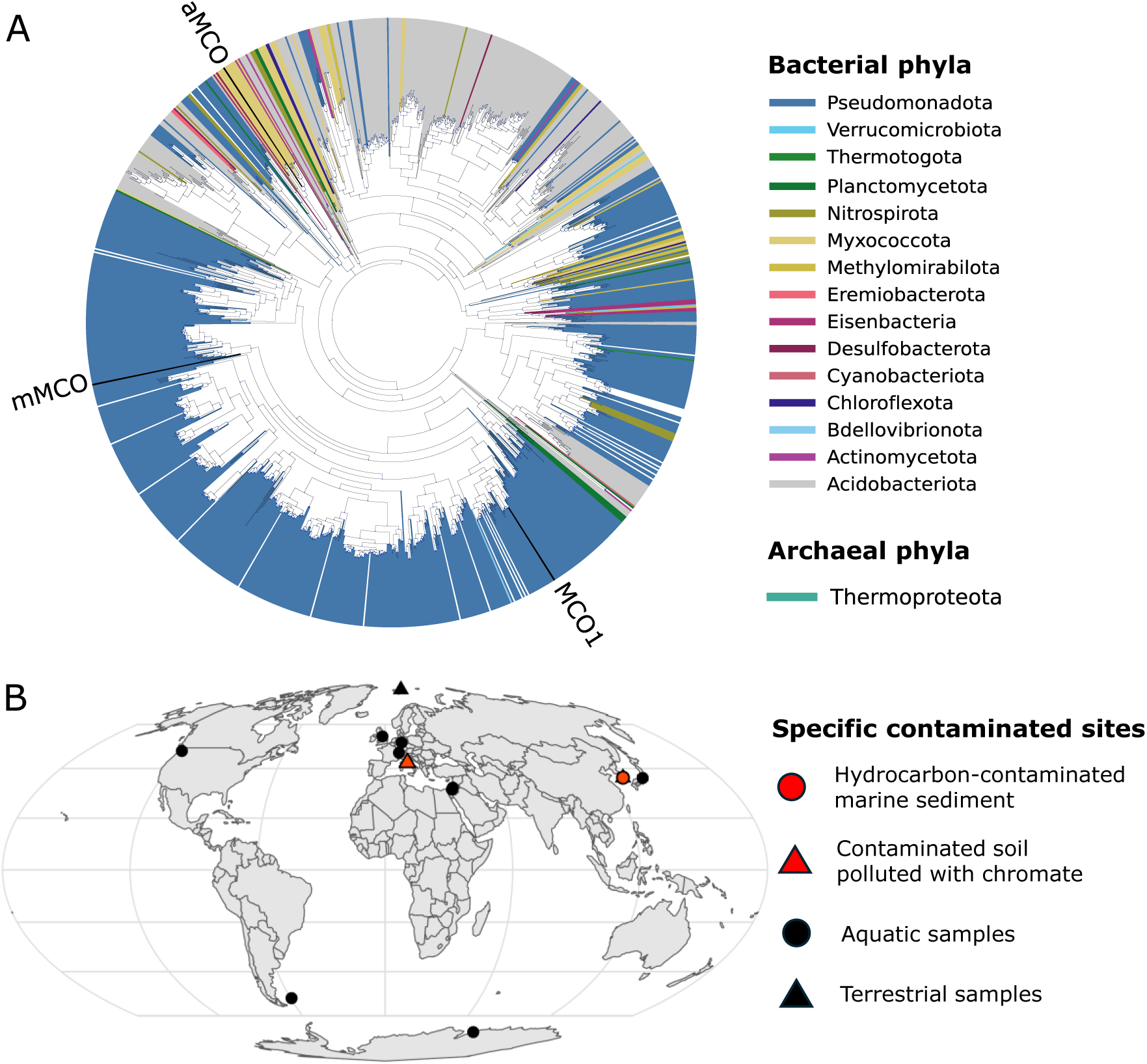
(A) Approximate maximum-likelihood phylogenetic tree of the multicopper oxidase MCO1 from *Nitrosomonas* sp. and two new laccases experimentally validated in this study from a methane-oxidizing bacterium (mMCO, NCBI ID: HEY7867587.1) and a soil ammonia-oxidizing archaeon (aMCO, HYV56308.1) displayed relative to other uncharacterized homologs in global metagenomic datasets colored by metagenomically-assigned phylum of origin. (B) Biogeographic distribution of MCO1 homologs from samples where location information was recorded in environmental sample metadata. Two samples were from notably contaminated sites: Red triangle = soil site contaminated with 1000 mg Cr (VI) kg^-1^. Red circle = crude-oil contaminated marine sediment site.

We identified MCO1 homologs from seventeen different bacterial phyla and one archaeal phylum (Figure 1). Among these, MCO1 homologs were detected in the major genera of ammonia-oxidizing microorganisms including *Nitrosomonas, Nitrosospira,* and the ammonia-oxidizing archaeon Ca. *Nitrosopolaris* sp. (HYV56308.1, phylum *Thermoproteota*, classified in NCBI as the *Nitrososphaerota* and formerly known as the *Thaumarchaeota*^38^). MCO1 homologs were also detected in major methane-oxidizing genera including *Methylobacter, Methylocystis, Methyloglobulus, Methylosarcina*, and numerous taxa from the phylum *Methylomirabilota* (formerly NC10). Apart from MCO1, however, none of these bacterial or archaeal-derived laccases have activity which has been determined experimentally. Among the 1103 curated MCO1 homologs identified in our analysis (Table S1), we therefore selected two additional candidates to evaluate their oxidation potential (enzyme sequences are noted in Table S1). These candidates were selected on the basis of being divergent MCO1 homologs belonging to different phyla than the original MCO1 (*Nitrosomonas*, belonging to the phylum *Pseudomonadota*). Namely, the candidates belonged to phyla known for their methane oxidation potential (*Methylomirabilota*, here termed mMCO) and an archaeal phylum with ammonia-oxidizing members (*Thermoproteota*, here aMCO). The genes encoding these enzymes were synthesized, cloned, heterologously expressed, and purified (Figure S1). We tested for oxidation activity with the model laccase substrate ABTS (2,2′-azino-bis(3-ethylbenzothiazoline-6-sulfonic acid), a commonly used redox mediator. Both mMCO and aMCO exhibited oxidation activity with ABTS (Figure S1). The ability of these MCO1 homologs to oxidize ABTS indicates that oxidative capability is conserved, even within a divergent archaeal phylum.

### High-throughput screening of 183 TrOCs biotransformation in a miniaturized laccase–mediator system

Having established that MCO1 is widely distributed in the environment and its function is conserved across diverse microbial lineages, we next used this enzyme to explore its reactivity against a broader chemical space. To create an enzyme-level benchmark suitable for predictive modeling, we assembled a library of 183 wastewater-relevant trace organic contaminants and quantified their biotransformation by MCO1 under standardized conditions.

To enable higher throughput screening while maintaining a chemically controlled oxidative environment, we implemented a miniaturized laccase–mediator system (LMS) using a 96-well plate setup and ABTS as the model mediator (Figure 2A)^6, 26^. Across all compounds, 38 exhibited significant biotransformation (criteria detailed in Methods) (Figure 2B, Table S2). Several compounds that showed extensive biotransformation in previous larger-volume, higher-enzyme-load assays displayed lower reactivity here (Table S2). For instance, amisulpride, sulfadiazine, and isoproturon, which exhibited more than 70% removal in the previous investigation, did not meet the significant biotransformation criteria in this scaled-down high-throughput assay (Table S2)^26^. Ranitidine, which was fully degraded in the high-enzyme-load system, demonstrated only partial removal under the current conditions (Figure 2C). Nonetheless, the significant biotransformation observed for 38 TrOCs (Figure 2B) demonstrates that this miniaturized LMS is well suited for surveying enzyme-substrate reactivity across large and diverse contaminant libraries.

**Figure 2.**
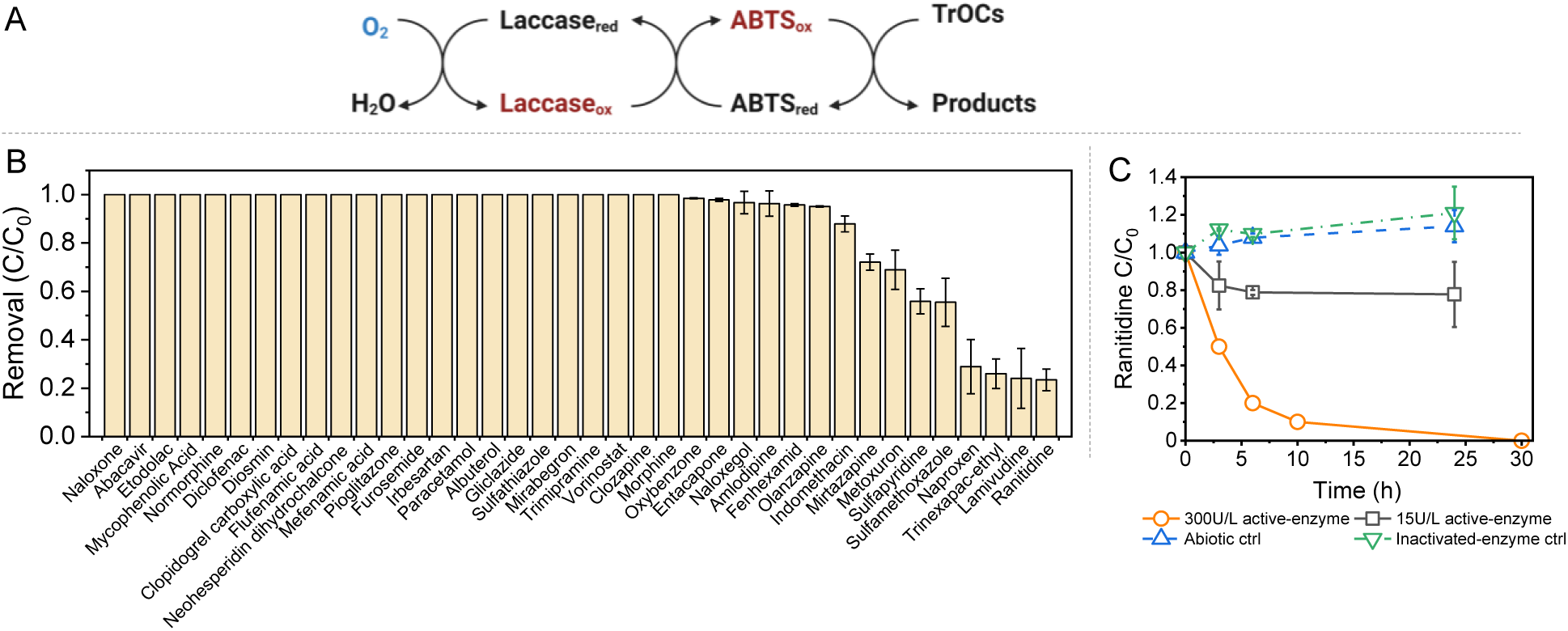
Biotransformation of TrOCs in the LMS. (A) Schematic of the laccase-mediator system; (B) TrOCs removal in the 96-well high-throughput screening assay (15U/L active enzyme in a 250 μL reaction volume); (C) Comparison of reactivity for ranitidine in different reaction systems (yellow, circle: 300U/L active enzyme in a 1000 μL reaction volume; black, box: 15U/L active enzyme in a 250 μL reaction volume; blue, up-triangle: abiotic buffer control with 250 μL reaction volume; green down-triangle: inactivated enzyme control with 250 μL reaction volume).

Compared to 24-well plate setups that are better suited for deriving robust kinetic rate constants under environmentally relevant conditions^39^, we used a 96-well format to enable parallel screening of 183 contaminants distributed across 14 sub-mixtures (including all controls) under a unified reaction framework. To balance this focus on throughput with the need for downstream model input, we propose a refined data-interpretation framework (Figure S2) beyond a traditional binary classification of “reactive” versus “non-reactive.” Specifically, we define a third “inconclusive” category to capture compounds whose apparent reactivity is sensitive to assay configuration or falls within low-signal fluctuation ranges (Figure S2). Compounds are thus assigned to “reactive”, “non-reactive”, or “inconclusive”, where the “inconclusive” group represents borderline behaviors that cannot be distinguished reliably under the current conditions. We exclude these “inconclusive” cases from the initial predictive modeling to reduce bias. Collectively, the miniaturized LMS provides a scalable platform for profiling enzyme–substrate interactions.

### Graph Convolutional Network (GCN) model for TrOCs biotransformation prediction

To evaluate whether enzyme-level reactivity data can support predictive modeling, we trained machine-learning classifiers on the curated LMS dataset. After excluding compounds categorized as “inconclusive”, 138 chemicals with confident “reactive” (n=38) or “non-reactive” (n=100) assignments were retained for model development. We first applied GCNs to construct a model for chemical biotransformation prediction. GCNs are particularly suitable for predicting biotransformation potential due to their ability to naturally incorporate the graph-like structure of chemical compounds into their learning process. By leveraging the graph representation of molecules, where atoms are nodes and bonds are edges, GCNs can effectively learn complex patterns and interactions crucial for distinguishing between reactive and non-reactive compounds. Unlike traditional molecular descriptors, which rely on pre-defined features and can miss subtle structural variations, GCNs automatically learn molecular features during training. To prevent overfitting, we first pre-trained our GCN using a contrastive learning approach on a dataset of 1 million PubChem molecules. Contrastive learning is a technique that helps the model learn meaningful representations by contrasting similar pairs and dissimilar pairs, encouraging the model to map similar molecules closer in the feature space and dissimilar ones further apart^40^. By doing so, the generalization capabilities of models can be improved when applied to downstream tasks, such as molecule classification or property prediction. After pre-training, an average classification accuracy of 0.78 ± 0.08 was achieved for the GCN model fine-tuned on 138 compounds (Table S2) with a 10-fold train-test split. By contrast, a benchmark multilayer perceptron (MLP) model trained on the same dataset using Molecular ACCess System (MACCS) descriptors as input achieved an average test accuracy of 0.73 ± 0.12 (Table 1).

**Table 1:** GCN prediction performance on train and test data. F1 is defined as the harmonic mean of precision and recall, providing a balanced measure of classification performance by jointly considering false positives and false negatives. AUROC represents the area under the receiver operating characteristic curve and reflects the model’s ability to discriminate between positive and negative classes across different classification thresholds.

|  | GCN |  | MLP with MACCS |  |
| --- | --- | --- | --- | --- |
|  | Train | Test | Train | Test |
| Accuracy | 0.84±0.02 | 0.78±0.08 | 0.76±0.01 | 0.73±0.12 |
| F1 Score | 0.65±0.06 | 0.63±0.10 | 0.58±0.02 | 0.51±0.08 |
| AUROC Score | 0.91±0.01 | 0.82±0.07 | 0.86±0.02 | 0.80±0.08 |

Furthermore, the 64-dimensional descriptors extracted from the trained GCN model effectively differentiated compounds that underwent biotransformation from those that did not (Figure 3A). The Principal Component Analysis (PCA) plot of these 64-dimensional descriptors for all compounds clearly showed a separation between the two classes (Figure 3A), whereas the PCA of the 167-dimensional MACCS descriptors (Figure 3B) failed to show such a distinction. These results highlight the potential of GCN models in capturing complex molecular patterns relevant to biotransformation prediction. By learning high-quality molecular representations directly from the graph structure of compounds, GCNs provide an excellent tool for predicting the biodegradability of new chemicals.

**Figure. 3.**
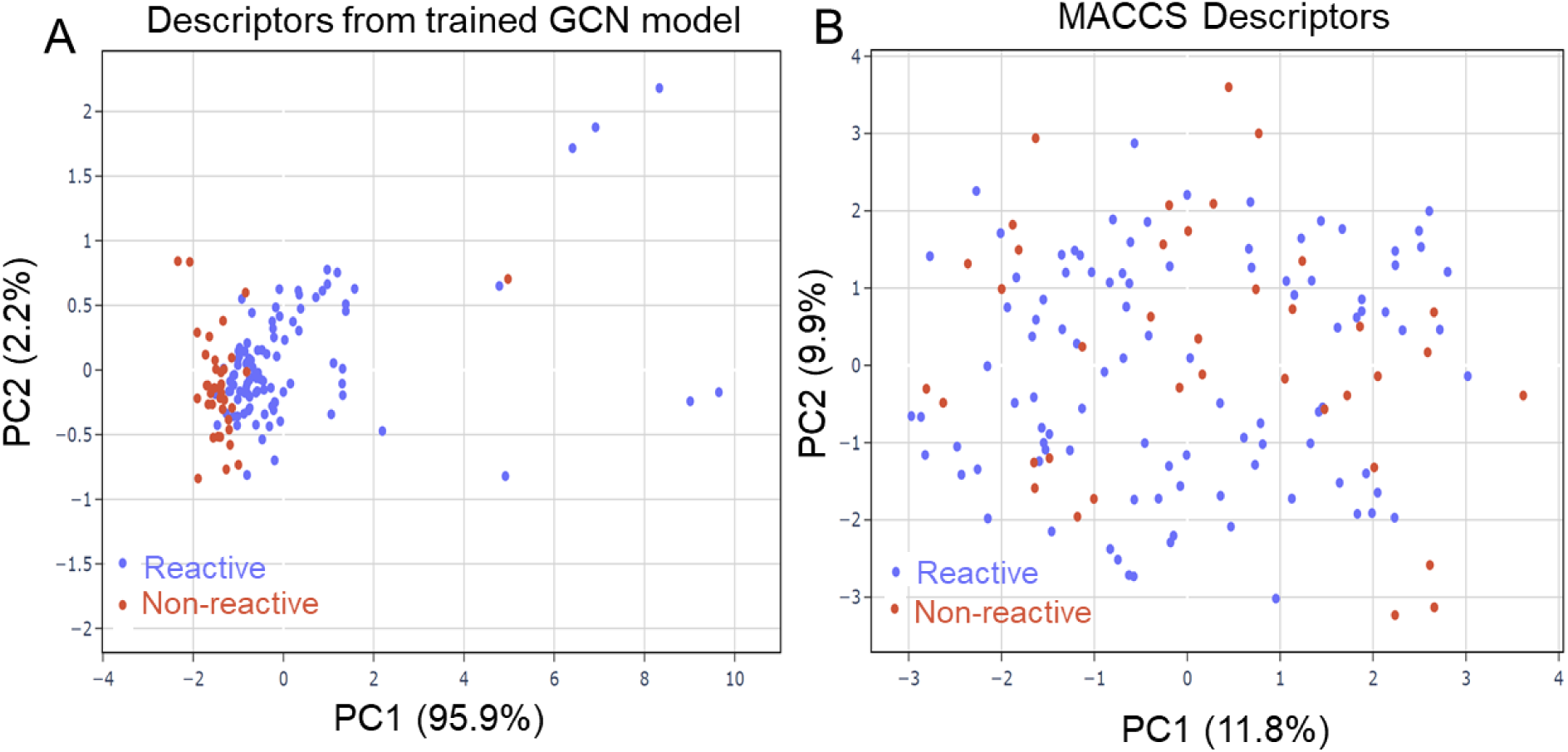
(A) Principal Component Analysis (PCA) of the descriptors generated by the 64-dimensional GCN model and (B) the MACCS fingerprints. Blue dots represent compounds with positive biotransformation, while red dots represent compounds with no biotransformation.

However, the descriptors generated by GCNs remain latent features, making it challenging to interpret the molecular factors driving individual predictions. For applications aimed at the development of SSbD chemicals, understanding why a molecule is predicted to be degradable or persistent, and which structural or chemical properties govern this outcome, is critical. Given that laccase-mediated oxidation is fundamentally governed by electron-transfer processes, we therefore explored whether quantum-chemical descriptors could complement the GCN framework by providing mechanistically-interpretable features. As a proof-of-concept, we analyzed a set of oxidation-relevant electronic parameters^41^ to evaluate whether they capture structure–reactivity relationships.

### Exploration of Descriptors from Quantum Chemical (QC) Calculations

Thanks to improved computational resources, it is now feasible to use quantities derived from advanced quantum chemical (QC) calculations as descriptors^41, 42^. The QC descriptors extend classic chemical descriptors based on electronic structure theory. In the case of the LMS, contaminant removal involves the interactions between TrOCs and the mediator-derived radicals, which ultimately initiate oxidation reactions. In other words, the key degradation process in the LMS is the chemical oxidation of the TrOCs by the mediator-derived radical. In the case of ABTS as mediator, the rate-limiting step is likely to be a one-electron transfer from the respective TrOC to the ABTS radical. Thus, we tried to find relevant QC parameters that can effectively capture the potential for TrOCs to be oxidized.

As an initial test, we examined the 20 molecules characterized in our previous large-volume experiment (LVE)^26^, including 16 molecules also present in the high-throughput LMS dataset, to directly compare between the two experimental setups. In order to search for indicators that can distinguish between the experimentally observed reacted and not-reacted molecular groups, we analyzed the correlations between the reactivity (degradability) and nine QC parameters: one-electron oxidation potential (*E_ox_*)^43^, electronegativity^44^, ionization energy^44^, electron affinity^44^, chemical hardness^44^, highest occupied molecular orbital (HOMO) energy, lowest unoccupied molecular orbital (LUMO) energy, HOMO-LUMO energy gap (*ΔE_HOMO-LUMO_*) and polarizability, calculated at the M062X/6-311+G(2df,2p)//M06L/6-311+G(2df,2p) level with the SMD solvent model^45^ (1^st^ QC). The results show that the combination of *E_ox_* and *ΔE_HOMO-LUMO_* clearly distinguishes between reacted and not-reacted molecules in LVE (Figure 4A). The classification was further validated by employing a support vector machine (SVM) classifier, achieving a cross-validation accuracy score of 0.95, a near-perfect accuracy. Importantly, these two parameters are fundamentally related to oxidation potential, providing a solid theoretical basis for separating the two classes of molecule.

**Figure 4.**
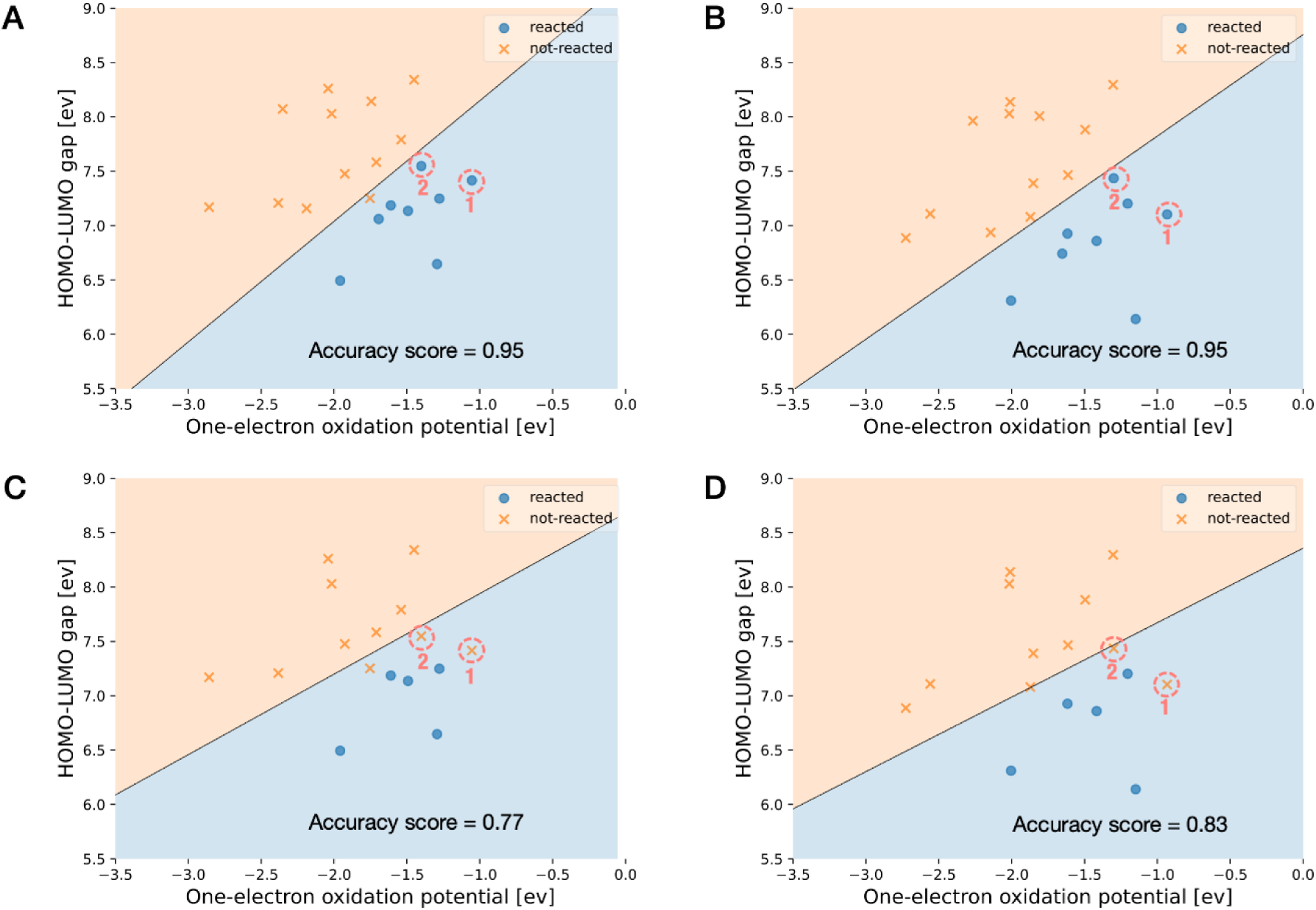
SVM classification of molecules used in the large-volume experiment (LVE) (20 molecules)^26^ and in this study (16 molecules). Amisulpride (1) and isoproturon (2) were reactive in LVE but not-reacted in this study. (A) LVE data for 20 molecules calculated at the 1^st^-QC level. SVM weight (the number of reacted compounds: the number of not-reacted compounds) = 1:1. (B) LVE, 20 molecules, 2^nd^-QC, SVM weight = 1:1. (C) This study, 16 molecules, 1^st^-QC, SVM weight = 0.45:1. (D) This study, 16 molecules, 2^nd^-QC, SVM weight = 0.45:1.

Encouraged by these promising results, we extended this approach to the 138 molecules with confident reacted or not-reacted assignments in the curated LMS dataset. In order to effectively handle the larger set of compounds, we opted for a slightly lower level in the QC calculations (2nd-QC) allowing for accelerated calculations while keeping the necessary accuracy for the relevant parameters (see Materials and Methods). The 2^nd^-QC level yields the same classification accuracy of 0.95 for the reactivity in LVE (Figure 4B), which demonstrated its suitability.

We used the 16 molecules used in both experimental systems for a first test. Under the miniaturized LMS conditions in this study, two compounds (1: amisulpride and 2: isoproturon) were newly determined to be non-reactive due to the lower activity of the miniatured LMS as discussed before. These discrepancies reduced the classification accuracy to 0.77 (1^st^-QC) and 0.83 (2^nd^-QC) (Figure 4C–D). When the analysis was extended to the 138 molecules, the boundary between the two classes became less distinct with a decreased accuracy of 0.71 (Figure 5A). The reacted molecules are distributed in the bottom right region, while the top left region contains mostly unreacted molecules. Although the separation is not perfect, this model can still predict rather confidently that an entry in the upper left region is a non-reactive molecule. The original QC calculations treated molecules in their neutral forms without consideration of ionization states. However, several compounds are expected to exist predominantly in ionic rather than neutral forms at the experimental pH of approximately 5–5.5. We then selected 58 from 138 molecules, which were predicted to be predominantly neutral at pH 5 by the Chemaxon method^46^. However, the results did not improve, showing a slightly lower SVM accuracy of 0.67 (Figure 5B). The results indicate that while these two QC descriptors remain important, a more general predictive model will likely require additional molecular descriptors to capture other confounding factors not represented by the current electronic features.

**Figure 5.**
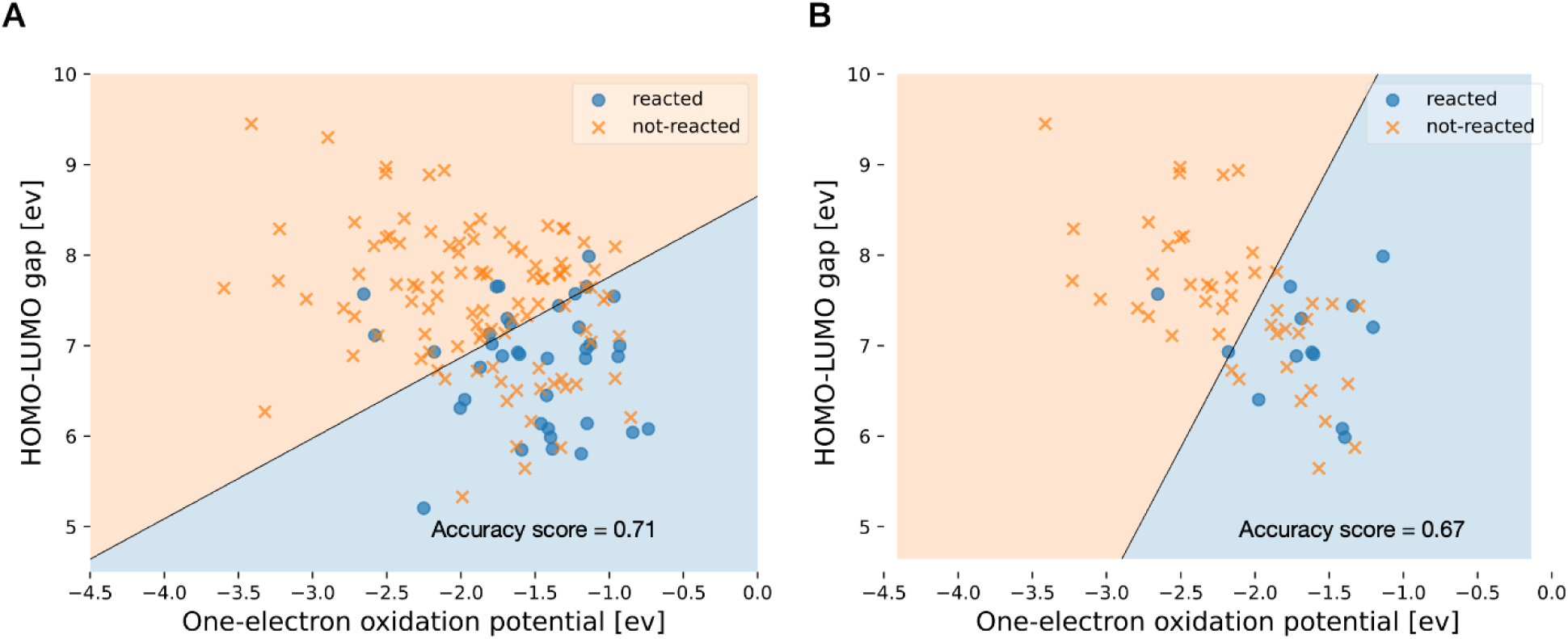
SVM classification of compounds in this study. (A) 138 compounds at the 2^nd^-QC level, SVM weight = 0.4:1. (B) 58 neutral molecules, 2^nd^-QC, SVM weight = 0.3:1.

### Environmental Implications

Predicting differences in biotransformation of TrOCs across environmental systems remains a fundamental challenge. Existing approaches rely on parameters such as pH, oxygen demand or organic carbon content, which reflect system conditions but do not capture the microbial processes that ultimately determine whether a chemical transforms or persists. Although the need to incorporate microbial signatures into predictive frameworks has been well recognized, progress has been limited because metagenomic and transcriptomic datasets are complex, redundant and only loosely connected to actual chemical reactivity. These limitations highlight the importance of furthering our understanding of chemical-enzyme interactions and developing prediction models that originate from chemical–enzyme interactions rather than from indirect community-level descriptors.

This study demonstrates an enzyme-level benchmark for bacterial laccases belonging to the MCO1-family. It additionally broadened the known sequence space of MCO1-family homologs by purifying and testing the activity of laccases from a newly characterized methane-oxidizing bacteria phylum and an ammonia-oxidizing archaeal phylum. Historically, the contribution of ammonia-oxidizing microorganisms to TrOC biotransformation has been attributed to the catalytic promiscuity of ammonia monooxygenase (AMO)^47, 48^, a copper-dependent membrane-bound enzyme known for its broad substrate specificity. However, validating the direct role of AMO in TrOC biotransformation has been challenging because such membrane-bound enzymes cannot be readily tested *in vitro*^49^. Evidence from previous studies already hinted that AMO is unlikely to be the only contributor^50^. For example, in nitrifying activated sludge, inhibition with the copper chelator allylthiourea caused a stronger reduction in TrOC biotransformation than treatments with more AMO-specific inhibitors such as octyne^50^. This pattern suggests that additional copper-dependent enzymes within nitrifying communities contribute to TrOC removal. Our findings support this broader view, suggesting a previously overlooked role of bacterial and archaeal laccases in contaminants removal.

We screened the activity of the ammonia-oxidizing bacterial laccase MCO1 across a set of >180 wastewater-relevant TrOCs. We combine this high-throughput enzymatic dataset with machine learning and quantum-chemical descriptors to derive mechanistically grounded predictions of biotransformation potential. This work represents an initial step toward building predictive tools that link chemical structure, enzyme specificity and reaction thermodynamics. Although this is a pilot study, the workflow is inherently scalable, and the integration of experimental enzymatic data with mechanistic descriptors offers a versatile foundation for future model expansion and multi-enzyme extensions. One potential direction is to integrate the QC descriptors with conventional QSAR features, which may better capture both local reactivity and global molecular behavior, thereby improving robustness across diverse chemical libraries. In addition, when more experimental records become available in large and chemically diverse datasets, more flexible modeling approaches, including deep learning architectures, could be explored to learn complex interactions between QC descriptors and chemical endpoints.

To evaluate how far this bottom-up framework can extend toward describing behavior in a relevant microbial community, we compared our results with an activated-sludge study that examined the same set of chemicals^51^. Ten of the forty-two compounds classified as undergoing oxidative catabolic degradation in that study were also transformed in our enzyme assay (Table S2), representing approximately one quarter of the catabolic group. At least another six compounds active in our assay showed behavior that made a clear delineation between catabolic and co-metabolic transformation difficult in the activated sludge study. This overlap is notable given that our system contains only one oxidative enzyme yet still seems to capture a measurable fraction of community-level oxidative biotransformation. As additional enzyme families are incorporated, this approach is expected to expand in predictive coverage and provide a pathway toward mechanism-informed tools that support environmental fate assessment and the design of more degradable chemicals.

## Materials and Methods

### Bioinformatic analysis

MCO1 homologs in different publicly available sequencing databases were filtered for protein sequences meeting the following criteria: at least 35% amino acid sequence identity, at least 85% alignment, and a bit score greater than 270. The National Center for Biotechnology Information (NCBI) non-redundant protein sequences database (nr) was queried using BLASTP^52^ with the above-listed stringent homology parameters, resulting in 960 unique protein sequences. Since the nr database is not comprehensive for environmental metagenomic sequences, we additionally queried three other databases with the same criteria: the Global Microbial Gene Catalog^27^, the BV-BRC database^28^, and the Joint Genome Institute Integrated Microbial Genomes (JGI-IMG) database^29^. All databases were accessed on 14 June, 2024. All hits were collated and duplicates were removed, resulting in 1,103 non-redundant MCO1 homologs.

Protein sequences were aligned using the DECIPHER sequence- and structure-based multi-sequence alignment algorithm^53^ The protein alignment was assessed and alignment edges (first 255 alignment positions, last 2078 positions) were trimmed. The trimmed alignment was used for approximate maximum likelihood phylogenetic tree estimation using FastTree v2.0^54^. Tree clades were colored based on phylum information from Genome Taxonomy Database release version 220 (GTDB)^55^, using the ETE 3 toolkit v 3.1.3^56^. For the BV-BRC sequences with linked metadata, geographic coordinates were extracted and mapped using rnaturalearth^57^.

### Heterologous gene expression

The investigated multicopper oxidase encoding gene (i.e., MCO1) was identified from the metatranscriptomics sequencing dataset of activated sludge samples, as previously described^26^. For gene expression, the MCO1-transformed *Escherichia coli* BL21(DE3) cryo stocks (stored at –80°C) were streaked onto LB agar plates supplemented with 100 μg/mL ampicillin and incubated at room temperature for 48 h. A single MCO1-expressing colony was picked and inoculated into 5 mL of terrific broth containing 100 μg/mL ampicillin and cultured overnight at 37°C with shaking. A 500 μL aliquot of this pre-grown culture was then transferred into 500 mL of terrific broth supplemented with 100 μg/mL ampicillin and incubated at 37°C with shaking at 200 rpm until the optical density at 600 nm (OD_600_) reached approximately 0.5. Protein expression was induced by adding 1 mM isopropyl-β-D-thiogalactopyranoside (IPTG). To promote the formation of the functional multicopper oxidase, CuCl_2_ was added together with IPTG at a final concentration of 0.25 mM. Cultures were transferred to 18°C and shaken for 4 h, followed by overnight incubation at 18°C without shaking to create microaerobic conditions favorable for active enzyme production. Cells were harvested by centrifugation at 5,000 × *g* for 10 min, and the cell pellets were stored at −20°C until use. For the two selected MCO1 homologs, protein expression was carried out using *Escherichia coli* T7. All subsequent induction and cultivation steps followed the same procedures described above for MCO1.

### Enzyme purification and SDS page

The cell pellets were resuspended in lysis buffer on ice (50 mM Tris-HCl, pH 7.5, 300 mM NaCl, 50% glycerol, 5 mM imidazole, 1 mM PMSF, 1 mg/mL lysozyme) and lysed using a French press (Avestin Emulsiflex C3). Cell debris and other impurities was removed by centrifugation, and the resulting supernatants were collected for enzyme purification. For benchtop purification, supernatants were applied to a Ni-NTA agarose column (Qiagen) pre-equilibrated with buffer A (20 mM Tris-HCl, 500 mM NaCl, 10% v/v glycerol, and 20 mM imidazole). The column was then washed with buffer A followed by two additional washes with buffer B (40 mM Tris-HCl, 500 mM NaCl, and 10% v/v glycerol). Final proteins were eluted using buffer C (500 mM imidazole, 500 mM NaCl, and 10% v/v glycerol). Protein fractions were collected and analyzed by SDS-PAGE (Figure S1A and Figure S1B). Fractions containing enzyme of interest, consistent with their respective molecular weights, were pooled and subjected to buffer exchange into TNG buffer (50 mM Tris-HCl, pH 7.4, 0.1 M NaCl, and 10% v/v glycerol) to remove imidazole. Purified enzymes were aliquoted, flash-frozen in liquid nitrogen, and stored at −80°C until further use. For the purification of MCO1 homologs, Fast Protein Liquid Chromatography (FPLC) was performed using an ÄKTA Pure system (Cytiva) following the procedures established in our previous work. After buffer exchange into TNG buffer, the purified homologs were aliquoted and stored at −80°C as described above for MCO1.

### Screening of laccase activity on a 96-well plate using a microplate reader

Laccase activity was quantified colorimetrically using 2,2’-azino-bis(3-ethylbenzothiazoline-6-sulfonic acid) (ABTS) as a chromogenic substrate (Figure S1C and Figure S1D). The assay followed a modified protocol adapted from previously described methods^26^. For MCO1, enzyme concentrations were first determined using the Bradford colorimetric assay. For activity measurements, enzyme stock solutions were diluted in ammonium acetate buffer (pH 5.6). Reactions were carried out in clear, flat-bottom 96-well plates by mixing 20 μL of diluted enzyme samples with 180 μL of buffer containing 500 μM ABTS. Laccase activity was quantified based on the initial rate of absorbance increase at 420 nm within 5 seconds of shaking at 30°C in a microplate reader. Kinetic measurements were conducted for one hour at one-minute intervals. The laccase activity toward ABTS is determined based on Equation 1^58^. For the two additional MCO1 homologs, enzyme activity was assessed qualitatively after purification by directly adding the enzyme to ABTS solution and visually confirming oxidation based on the characteristic color change (Figure S1D).

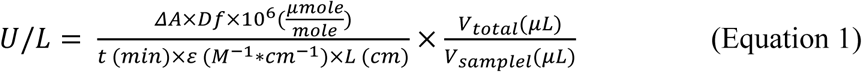

Where, **U**: enzyme activity (μmole/(min*L)); **ΔA**: the change in absorbance and is calculated as the final absorbance minus the initial absorbance; **Df**: dilution factor; **t**: reaction time (min); **ɛ**: molar extinction coefficient (36000 M^-1^ cm^-1^, M=mole/L); **L**: path length, which is 0.56 cm for a 96-well plate containing 200μL of solution; **V_total_**: total volume of the reaction (μL); **V_sample_**: Sample volume (μL).

### Compound selection and clustering

A total of 183 wastewater-relevant trace organic contaminants (TrOCs), including commonly used pesticides, pharmaceuticals, and artificial sweeteners, were selected for this study. Individual stock solutions were prepared in dimethyl sulfoxide (DMSO) at 10 mM and stored at −20 °C until use. To reduce potential competitive inhibition in the high-throughput screening assay, we partitioned the 183 compounds into 14 sub-mixtures (final concentration at 200 μM for each TrOC in ethanol) using hierarchical clustering. The clustering was based on the compound structural information together with predicted biotransformation reaction rules from enviPath^3, 59^. This design distributes structurally and mechanistically similar compounds across different sub-mixtures, thereby limiting the number of closely related substrates co-incubated in the same well and reducing the likelihood of strong mixture effects (*e.g.*, shared-enzyme competition or combined toxicity). All sub-mixtures were stored at −20 °C for subsequent biotransformation experiments. Detailed sub-mixture information is included in Table S2, same clustering strategy has been reported previously^6^.

### High-throughput biotransformation assay of the investigated enzyme and selected TrOCs

TrOCs biotransformation by MCO1 was assessed using a clear-bottomed 96-well plate (Greiner). Individual TrOC sub-mixtures were first added to an empty 96-well plate. After complete evaporation of the organic solvent (i.e., ethanol), 240 μL of ammonium acetate buffer (pH ∼5.5) was added to each well. The plate was then subjected to shaking at 50 rpm for 15 minutes to redissolve the TrOCs. Subsequently, 5 μL of ABTS stock solution (final concentration 200 μM) was added to the well, followed by the addition of 5 μL of purified enzyme solution (with a final enzyme activity of ∼15 U/L). The plate was incubated in a self-designed humidified chamber at 30°C^6^, with continuous shaking at 150 rpm. To monitor potential evaporation, the total weight of the plate was measured before and after each sampling event. No significant evaporation was detected during the experimental period. Additionally, to ensure sufficient oxygen exposure for the enzyme, in addition to shaking, each well in the 96-well plate was aerated thoroughly using a multichannel pipette every three hours throughout the duration of the experiment. Samples for LC-HRMS/MS analysis were collected at 0, 3, 6, and 24 hours. At each sampling event, a 20 μL sample was taken and mixed with the same amount of pure methanol to completely denature the enzyme. A 160 μL of ammonium acetate buffer was then added to the sample mixture to achieve an approximate 10-fold dilution of the original sample. Following centrifugation at 13,000 × *g* for 15 minutes at 4°C, the supernatant was collected and stored at 4°C for LC-HRMS/MS analysis.

Additionally, enzyme-free abiotic fresh buffer control and trichloroacetic acid (TCA) treated inactivated enzyme control were prepared in parallel with the biological experiment. All experiments were conducted in duplicate, and samples were measured by LC-HRMS/MS within two days of collection. Matrix-matched calibration standards were used for quantification, with concentrations determined from peak area integration using TraceFinder 5.1 (Thermo Fisher Scientific).

### Ultrahigh-performance liquid chromatography coupled to high-resolution tandem mass spectrometry (UHPLC-HRMS/MS) analysis

The TrOCs were analyzed using UHPLC–HRMS/MS on a Q Exactive system (Thermo Fisher Scientific) following a method adapted from our previous work^6^. Briefly, for the LC separation, 25 μL of each sample was loaded onto an ACQUITY Premier BEH C_18_ column (1.7 μm, 100 × 2.1 mm, Waters) and eluted with nano-pure water (A) and methanol (B), each containing 0.1% formic acid. The flow rate was set at 300 μL/min. A linear elution gradient was as follows: 95% A: 0 – 1.5 min, 95% – 5% A:1 – 7.5 min, 5% A: 7.5 – 9.5 min, and 95% A: .5 – 11.5 min. For HRMS, mass spectra were acquired in full scan mode with a resolution of 70,000 at m/z 200, using positive/negative switching electrospray ionization mode over a scan range of m/z 100 – 1000. Xcalibur 4.0 (Thermo Fisher Scientific) and TraceFinder 5.1 (Thermo Fisher Scientific) were used for data acquisition and analysis. ChemDraw Professional 20.0 and MarvinSketch (v19.20.0, ChemAxon, http://www.chemaxon.com) were used for drawing, displaying, and characterizing chemical structures

### Data Cleaning Framework for Predictive Modeling in Chemical Fate and Biotransformation Studies

Current predictive models for TrOC biotransformation or chemical fate assessment are often built from datasets curated from the literature, where compound classifications typically rely on strict but somewhat arbitrary criteria. A commonly used approach defines “Significant Biotransformation” as >20% biological removal relative to abiotic and dead-cell controls (Figure S2)^47^. Although such thresholds help minimize false positives, they may also misclassify weakly reactive or slowly transforming compounds as non-reactive, thereby biasing predictive models toward reduced sensitivity for these compounds. To better align experimental measurements with computational model requirements, here, we applied a refined classification scheme that introduces an intermediate “inconclusive” category (Figure S2).

In the dataset used for modeling (Table S2), the maximum removal percentage of individual compounds for different control groups was calculated using Equation 2. Compounds were classified as Positive if they exhibited more than 20% removal and significantly higher removal in the biological control compared to both abiotic and inactive-enzyme controls (*p* < 0.05), consistent with criteria used in previous biodegradation studies. Rather than grouping all remaining compounds as Negative, we assigned compounds showing 10–20% removal and/or lacking statistically significant differences from controls (p ≥ 0.05) to the “inconclusive” category (Figure S2). Compounds with less than 10% removal were classified as Negative, indicating their persistence under specific conditions. Compounds assigned to the “inconclusive” category were excluded from model training. This selection strategy ensures that the predictive model is calibrated with only reliable data, resulting in more accurate and robust predictions of chemical biotransformation. By applying this criterion, we identified 38 positive and 100 negative results suitable for downstream computational modeling. The detailed compounds classification is included in Table S2.

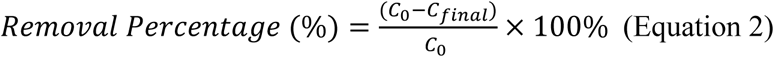

Where, **C_0_**represents the initial concentration measured by LC-HRMS/MS, and **C_final_**denotes the final concentration after biotransformation experiments.

### Graph Convolutional Network (GCN) model construction

Our Graph Convolutional Network (GCN) model, an extension of the neural network class (nn.Module) in PyTorch, was designed for graph-structured data analysis. The model was initialized with three graph convolutional layers to process the features of nodes, each followed by a ReLU activation function to introduce non-linearity. Convolution layers progressively transformed the node features from the input space to a hidden space comprising 64 channels. To aggregate features of nodes into a graph-level representation, we applied global mean pooling and achieved a 64-dimensional descriptor for each compound. Our model contained a dropout layer for regularization and a linear classifier that mapped the 64-dimensional descriptor to the class space of the dataset. In addition, we designed a dataset class for the systematic conversion of molecular data into graph representations, suitable as input for our model. This class processed molecules encoded as SMILES strings, translating them into graphs and extracting the adjacency information of atoms. Within these graphs, nodes represented atoms and edges denoted chemical bonds. The node features encompassed atomic number, degree, formal charge, hybridization, aromaticity, total hydrogen count, radical electron count, inclusion in rings, and chirality. The edge features included bond type and ring membership. Ten neutral compounds were randomly selected as the test set, while the remaining 128 compounds were used for training. The train-test split was repeated 10 times to ensure a comprehensive evaluation. We also developed a benchmark model by removing the convolutional components from our GCN model and retaining only the final linear classifier, using MACCS descriptors as the input. It allows us to compare the 64-dimensional descriptors learned by the GCN model with the predefined MACCS descriptors^60^.

### Exploration of Quantum Chemical Descriptors

All QC calculations were performed using the Gaussian 16 program package^61^. PubChemPy (ver. 1.0.4) was used to obtain geometries that were used as the initial structures for QC calculations. We used Scikit-learn (ver. 1.1.3) for the SVM modeling and Mlxtend (ver. 0.23.1) and Matplotlib (ver. 3.6.2) for the plotting. The ionization states of each compound at pH 4-6 were determined using the Chemaxon pKa calculator (https://chemaxon.com/calculators-and-predictors).

The initial structures were obtained from PubChem (https://pubchem.ncbi.nlm.nih.gov) automatically from the compound names. If there is no three-dimensional geometry available, the initial structure was manually built by using GaussView6^61^. The first step is geometry optimization for the initial structure at the M06L/6-311+G(2df,2p) level with the SMD solvent model to give result 1 (R1) followed by a single point calculation for the structure of R1 at the M062X/6-311+G(2df,2p) level to give result 2 (R2). The geometry of R1 is used for geometry optimization of the cation as an open shell doublet system at the M06L/6-311+G(2df,2p) level with the SMD solvent model to give result 3 (R3), followed by a single point calculation to give result 4 (R4). Using these results, one-electron oxidation potential, 𝐸_𝑜𝑥_, was calculated by Equation 3^43^.

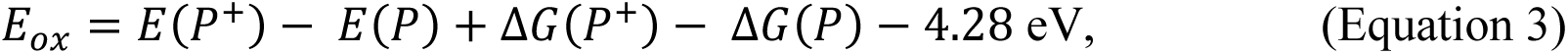

where, 𝐸(𝑃^+^) and 𝐸(𝑃) are the potential energies of R4 and R2, respectively, 𝑑𝐺(𝑃^+^) and 𝑑𝐺(𝑃) are thermal corrections to the Gibbs free energy of R3 and R1, respectively. The unit is electron volt (eV). The energies of anionic and cationic species for other parameters including electronegativity^44^ were obtained by single point calculations of open shell systems. HOMO and LUMO energies were obtained from R2.

To obtain the lower QC level for acceleration, we searched for a relevant level by comparing the accuracy of 𝐸_𝑜𝑥_ and Δ𝐸_𝐻𝑂𝑀𝑂−𝐿𝑈𝑀𝑂_ with respect to the 1^st^ QC level with the following four combinations of QC level and methods: (i) PM7//PM7, (ii) M062X/6-31+G(d,p) with SMD//PM7, (iii) M062X/6-31+G(d,p)//PM7 both with SMD and (iv) M062X/6-31+G(d,p)//BLYP/3-21G(d) both with SMD, where PM7 is a semi-empirical method, and SMD is a solvent model. The method before and after the notation “//” is for obtaining energy and geometry, respectively, i.e., the former was used for a single point calculation, and the latter was used for geometry optimization. We employed a dataset containing 167 molecules calculated at the 1^st^ QC level. The R^2^ value for 𝐸_𝑜𝑥_ and Δ𝐸_𝐻𝑂𝑀𝑂−𝐿𝑈𝑀𝑂_ calculated with these methods against those by the 1^st^ QC level are shown in Table S4. As shown in Table S4, most of the methods show a good correlation for Δ𝐸_𝐻𝑂𝑀𝑂−𝐿𝑈𝑀𝑂_ but mostly fail for 𝐸_𝑜𝑥_. Only method (iv) resulted in an acceptable R^2^ value (0.907). We inspected also the similarity of geometries as well as the relevancy of the parameter values as indicators to separate reactive and non-reactive molecules predicted by the SVM model built using the 20 molecules examined in the previous experimen^26^. Thus, we decided to use method (iv) as the 2^nd^ QC level. 𝐸_𝑜𝑥_ is calculated by Equation 3 but without thermal correlation to Gibbs free energy. Using this level, computation became about four times faster compared to that at the 1^st^ QC level.

### Training Support Vector Machine (SVM) Model

We use the linear kernel function for the SVM modeling. The cross-validation score was performed by five times splitting of data and fitting the model for the 20 and 16 molecules (Figure 4) and eight times for the 138 and 58 molecules (Figure 5). For plots in which “reacted” and “not-reacted” compounds are partly mixed, i.e., all plots with the current experiment (Figure 4C, Figure 4D, Figure 5A and Figure 5B), weights corresponding to the ratio of the number of “reacted” and “not-reacted” compounds was considered in the SVM modeling. The weight values are listed in the respective figure captions.

## Supporting information

Laccase_SI_tables_bioRxiv

Laccase_SI_bioRxiv_v5

## Supplementary information

The Supplementary Information includes four Supplementary Tables (Tables S1–S4), provided in an accompanying Excel file. Figures S1 and S2 present the SDS-PAGE characterization of purified MCO1 and its homologs, , as well as the proposed data-cleaning framework used for predictive modeling.

## CRediT Authorship Contribution Statement

**Yaochun Yu:** Conceptualization, Formal analysis, Investigation, Methodology, Visualization, Supervision, Writing – original draft, Writing – review & editing. **Kunyang Zhang:** Formal analysis, Investigation, Methodology, Writing – original draft, Writing – review & editing. **Vincenz-Maria Steiner:** Formal analysis, Methodology, Writing – review & editing. **Victoria Poltorak:** Formal analysis, Methodology, Visualization, Writing – review & editing. **Silke I. Probst**: Methodology, Writing – review & editing. **Serina L. Robinson:** Formal analysis, Investigation, Methodology, Visualization, Writing – original draft, Writing – review & editing. **Jürg Hutter:** Conceptualization, Supervision, Writing – review & editing. **Hiroko Satoh:** Conceptualization, Formal analysis, Investigation, Methodology, Visualization, Supervision, Writing – original draft, Writing – review & editing. **Kathrin Fenner:** Conceptualization, Supervision, Funding acquisition, Writing – original draft, Writing – review & editing.

## Data Availability

Scripts used for the bioinformatics analysis in this manuscript are available at : https://github.com/MSM-group/laccase-paper. Geometries of all compounds as well as all computational descriptors are available at https://github.com/shiroco/Data/tree/main/wasser-degradation/138. Scripts related to Graph Convolutional Network (GCN) model in this manuscript are available at https://github.com/zhangky12/mco1

## Acknowledgments

This work was supported by the Swiss National Science Foundation (project no. 200021L_201006, for Y.Y. and K.F.; PZPGP2_209124, no. 501100001711-209124 for V.P. and S.L.R.), the Eawag Academic Transition Grant (for Y.Y.), the European Union’s H2020 research and innovation program under the Marie Sklodowska Curie grant agreement (award no. MSCA-ITN-H2020[956496], for K.Z.), and the Peter und Traudl Engelhorn Stiftung (for S.I.P.). We thank Philipp Longree from the Department of Environmental Chemistry (Uchem) at Eawag for the method development on the UHPLC-HRMS/MS. We thank Elia Ceppi (Uchem, Eawag) for assisting with the sample measurement on UHPLC-HRMS/MS. We thank Cleo Soldini (Uchem, Eawag), Till Epprecht (Department of Environmental Microbiology, Umik, Eawag), Niklas Ferenc Trottmann (Umik, Eawag), and René Gall (Umik, Eawag) for their technical support on the enzyme purification. We thank Martina Kalt (Uchem, Eawag) and Miguel Heussi (Uchem, Eawag) for preparing chemical stock solutions. We thank Dr. Sema Karakurt-Fischer (Process Engineering Department, Eawag) for designing and kindly sharing the humidified chamber. We thank Dr. Anastasia Athanasakoglou (formerly a postdoctoral researcher at Uchem, Eawag) for the thorough discussions about the enzyme purification protocol.

