## Supplementary material for "An enzyme-level benchmark based on environmental bacterial laccases for predicting contaminant fate in water": Laccase_SI_bioRxiv_v5

### Table of contents

Supplementary Tables. All Supplementary Tables (S1–S4) are provided in the attached Excel file, and detailed descriptions of each table are listed below. ............................................................3
Table S1. MCO1 homolog search results used for constructing Figure 1. This table summarizes all MCO1 homologs identified in the sequence search and used for downstream phylogenetic analyses in Figure 1. ....................................................................................................................3
Table S2. Summary of biotransformation results. This table provides the detailed biotransformation results for all tested chemicals exposed to purified MCO1. The first column, *Overlap with Large-Volume Higher-Protein-Activity Control*, indicates the 16 compounds that overlapped with results obtained in our previous large-volume experiment. “Yes” denotes compounds that showed removal in the previous study, whereas “No” indicates no removal in previous study. The second column, *Overlap with Catabolic Biotransformation Compounds*, identifies chemicals in this dataset that overlap with the set of compounds previously classified as undergoing catabolic biotransformation in activated sludge. Additional biotransformation classifications (Reactive, Non-reactive, Inconclusive) are included to support downstream modeling analyses. .........................................................................................................3
Figure S1. Characterization of laccases. (A) SDS-PAGE analysis of MCO1. Protein molecular weight markers are shown in lanes 1 and 12. Lanes 2–11 contain cell debris samples, flow-throughs, and fractions collected throughout the purification workflow. The target protein MCO1 is clearly visible in the elution fractions in lanes 9 and 10. (B) Enzymatic activity of purified MCO1, measured by monitoring the oxidation of ABTS at 420 nm. (C) Colorimetric reaction of MCO1, aMCO, and mMCO with ABTS. Duplicate reactions with different enzyme volumes are shown, and the visible color change relative to the 0 µL enzyme control reflects the formation of the ABTS radical cation and confirms oxidative activity. ..................................................................................4
Figure. S2. Schematic flow diagram illustrating the data cleaning process for biotransformation experiments, comparing the traditional dataset approach (left) and the refined dataset (right) used for downstream computational modeling. .........................................................................5

**Supplementary Tables.** All Supplementary Tables (S1–S4) are provided in the attached Excel file, and detailed descriptions of each table are listed below.

**Table S1.** MCO1 homolog search results used for constructing Figure 1. This table summarizes all MCO1 homologs identified in the sequence search and used for downstream phylogenetic analyses in Figure 1.

**Table S2.** Summary of biotransformation results. This table provides the detailed biotransformation results for all tested chemicals exposed to purified MCO1. The first column, *Overlap with Large-Volume Higher-Protein-Activity Control*, indicates the 16 compounds that overlapped with results obtained in our previous large-volume experiment. "Yes" denotes compounds that showed removal in the previous study, whereas "No" indicates no removal in previous study. The second column, *Overlap with Catabolic Biotransformation Compounds*, identifies chemicals in this dataset that overlap with the set of compounds previously classified as undergoing catabolic biotransformation in activated sludge. Additional biotransformation classifications (Reactive, Non-reactive, Inconclusive) are included to support downstream modeling analyses.

**Table S3.** 58 neutral charge chemicals and their calculated one-electron oxidation potential and HOMO-LUMO gap.

**Table S4.** Accuracy and Efficiency Compared to the 1st QC level.

#### Supplementary Figures.

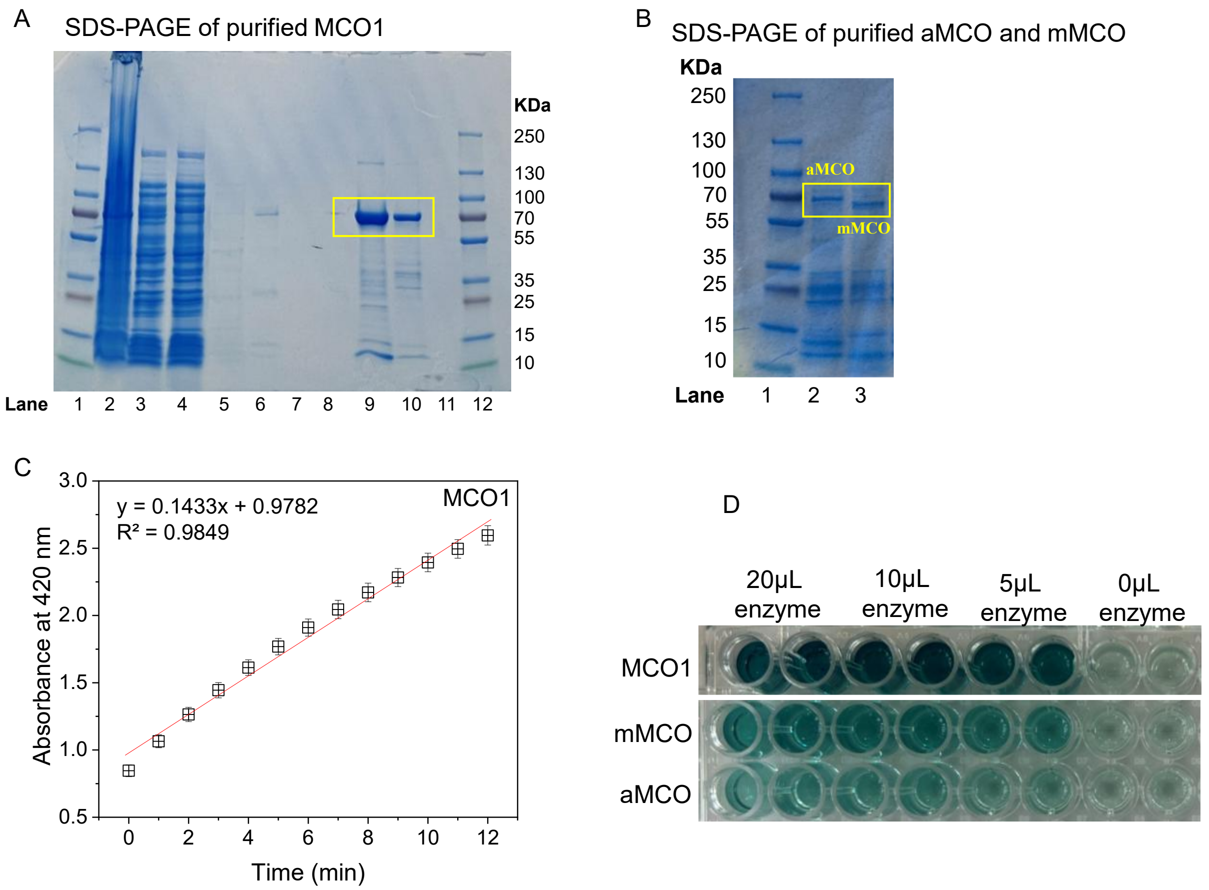

**Figure S1. Characterization of laccases.** (A) SDS-PAGE analysis of MCO1. Protein molecular weight markers are shown in lanes 1 and 12. Lanes 2–11 contain cell debris samples, flow-throughs, and fractions collected throughout the purification workflow. The target protein MCO1 is clearly visible in the elution fractions in lanes 9 and 10. (B) SDS-PAGE analysis of aMCO and mMCO (C) Enzymatic activity of purified MCO1, measured by monitoring the oxidation of ABTS at 420 nm. (D) Colorimetric reaction of MCO1, aMCO, and mMCO with ABTS. Duplicate reactions with different enzyme volumes are shown, and the visible color change relative to the 0 μL enzyme control reflects the formation of the ABTS radical cation and confirms oxidative activity.

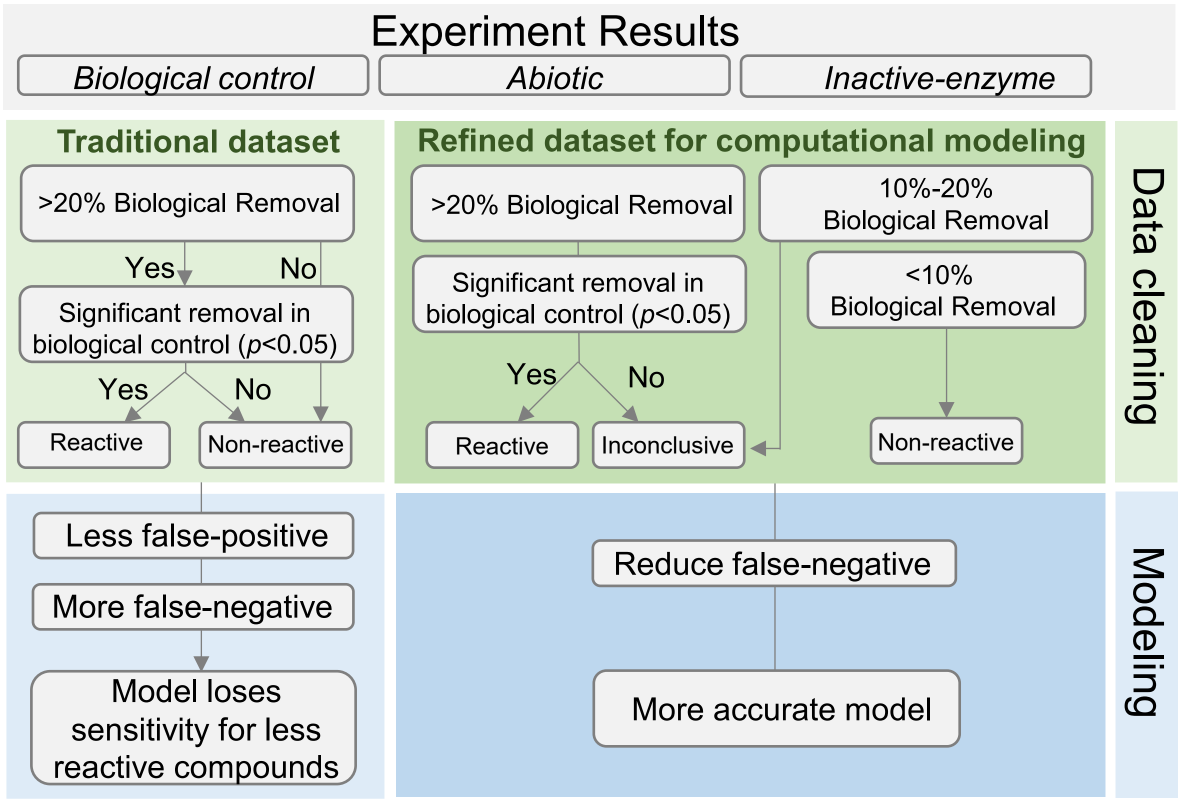

**Figure. S2.** Schematic flow diagram illustrating the data cleaning process for biotransformation experiments, comparing the traditional dataset approach (left) and the refined dataset (right) used for downstream computational modeling.
